# Screening of the optimal CpG-oligodeoxynucleotide for anti-inflammatory responses in avian macrophage cell line HD11

**DOI:** 10.1101/2022.04.05.487094

**Authors:** Kennosuke Ichikawa, Mei Matsuzaki, Ryo Ezaki, Hiroyuki Horiuchi, Yoshinari Yamamoto

**Affiliations:** Genome Editing Innovation Center, Hiroshima University, 3-10-23 Kagamiyama, Higashi-Hiroshima, Hiroshima 739-0046, Japan; Graduate School of Integrated Sciences for Life, Hiroshima University, 1-4-4 Kagamiyama, Higashi-Hiroshima, Hiroshima 739-8528, Japan

**Keywords:** anti-inflammation, chick macrophages, CpG-ODN, HD11, IL-10

## Abstract

CpG-oligodeoxynucleotides (**CpG-ODN**) has been shown to possess immunostimulatory features in both mammals and birds. However, compared to its pro-inflammatory effects, little is known about the anti-inflammatory responses triggered by CpG-ODN in avian cells. Hence, in this study, we aimed to characterize the anti-inflammatory response in the chick macrophage cell line HD11 under the stimulation of five kinds of CpG-ODN: CpG-A_1585_, CpG-A_D35_, CpG-B_1555_, CpG-B_K3_, and CpG-C_2395_. Single-stimulus CpG-B_1555_, CpG-B_K3_, and CpG-C_2395_ induced the interleukin (***IL****)-10* expression without causing cellular injury. The effects of pretreatment with each CpG-ODN before subsequent lipopolysaccharide stimulation were also evaluated. Interestingly, only CpG-C_2395_ maintained a high expression level in this situation. Finally, expression analysis of inflammation-related genes, such as the tumor necrosis factor*-*α, *IL-1β, IL-6*, and Toll-like receptor 4, was conducted, and pretreatment with CpG-C_2395_ significantly reduced their expression. Overall, our results shed light on the anti-inflammatory responses triggered by CpG-C_2395_ stimulation using a comparative analysis of three major classes of CpG-ODN in chick macrophages.

## INTRODUCTION

Macrophages contribute significantly to host immune responses via phagocytosis, antigen presentation, and cytokine secretion. Avian macrophages are dispersed throughout body fluids and tissues (Klasing, 1998) and are thus suited to regulate host immune responses in the whole body. Avian macrophages challenged with pathogens or immunogens express several pro-inflammatory cytokines, such as the tumor necrosis factor (**TNF**)-α, interleukin (**IL**)-1β, and IL-6, and induce immune and inflammatory responses (Klasing, 1994). However, the excessive expression of pro-inflammatory cytokines results in severe diseases. For example, septic shock, caused by the inappropriate regulation of immune responses, has a high mortality rate (Stearns-Kurosawa et al., 2011). Additionally, in birds, infection with the avian influenza virus inappropriately regulates the immune system (Burggraaf et al., 2014). Thus, the proper regulation of inflammatory responses in avian macrophages is critical to prevent immune system-related avian diseases in the poultry industry.

In this study, we focused on the immunostimulatory features of CpG-oligodeoxynucleotides (**CpG-ODN**) as potential regulators of immune responses. CpG-ODN directly upregulate cytokines in innate immune cells such as macrophages (Krieg, 2002). Several studies have demonstrated that CpG-ODN activate innate immune responses in chickens. For example, CpG-ODN induce inflammatory responses by upregulating nitric oxide, *IL-1β, IL-6*, and *IL-12* in a chicken macrophage-like cell line HD11 (He et al., 2003; He and Kogut, 2003; Xie et al., 2003; He et al., 2007; Han et al., 2010). Further, CpG-ODN not only induce pro-inflammatory responses but also upregulate IL-10, a significant factor for the anti-inflammatory response, in mammalian cells (Yi et al., 2002; Saraiva and O’Garra, 2010). However, knowledge about the anti-inflammatory response triggered by CpG-ODN in avian cells is scarce. Because the induction of IL-10 suppresses the excessive expression of pro-inflammatory cytokines, such basic knowledge can contribute to disease prevention in poultry.

CpG-oligodeoxynucleotides are divided into three classes —A, B, and C— based on their structural and biological characteristics (Krieg, 2006). Most of the studies that assessed the immunostimulatory features of CpG-ODN against chicken macrophages were conducted using Class B CpG-ODN (He et al., 2003; He and Kogut, 2003; Xie et al., 2003; He et al., 2007; Han et al., 2010). Although recent studies have used other classes of drugs (Ciraci and Lamont, 2011; Barjesteh et al., 2014), the anti-inflammatory response of chick macrophages against CpG-ODN has not been characterized.

Hence, in the present study, we compared cellular injury and *IL-10*-inducibility in HD11 cells stimulated with five different CpG-ODN belonging to each class. Here, two types of treatment, namely single-stimulus with CpG-ODN and pretreatment with CpG-ODN before subsequent lipopolysaccharide (**LPS**) stimulation, were conducted. During these experiments, we evaluated the anti-inflammatory responses triggered by CpG-C_2395_ and the effects of the pretreatment with CpG-C_2395_ before LPS stimulation for downregulating inflammation-related genes.

## MATERIALS AND METHODS

### Cell culture

The HD11 cells were propagated and maintained in complete Dulbecco’s modified Eagle’s medium (**DMEM**) (Thermo Fisher Scientific, Waltham, Massachusetts) supplemented with 10% (v/v) fetal calf serum at 37 °C with 5% CO_2_. HD11 cells were subcultured every 3 to 4 days. HD11 cells were seeded onto a 24-well cell culture plate (Corning Inc., New York) at a density of 1×10^5^ cells/well in complete DMEM and cultured for 24 h at 37 °C in a humidified incubator supplied with 5% CO_2_. After harvest, the cell culture supernatant was removed by aspiration, and the cells were washed once with PBS and used for further experiments.

### Treatments of HD11 cells with CpG-ODN and LPS

Endotoxin-free desalted phosphorothioate-ODN were synthesized by Gene Design Inc. (Osaka, Japan). Phosphorothioate-oligodeoxynucleotides were reconstituted in PBS and passed through a 0.22 µm pore microfilter (Nihon Millipore K.K., Tokyo, Japan). ODN sequences, namely CpG-A_1585_, CpG-A_D35_, CpG-B_1555_, CpG-B_K3_, CpG-C_2395_, and GpC_1612_, are shown in Table 1. Lipopolysaccharide from *Escherichia coli* O123:B8 was purchased from Sigma-Aldrich (St. Louis, Missouri).

**Table 1.**
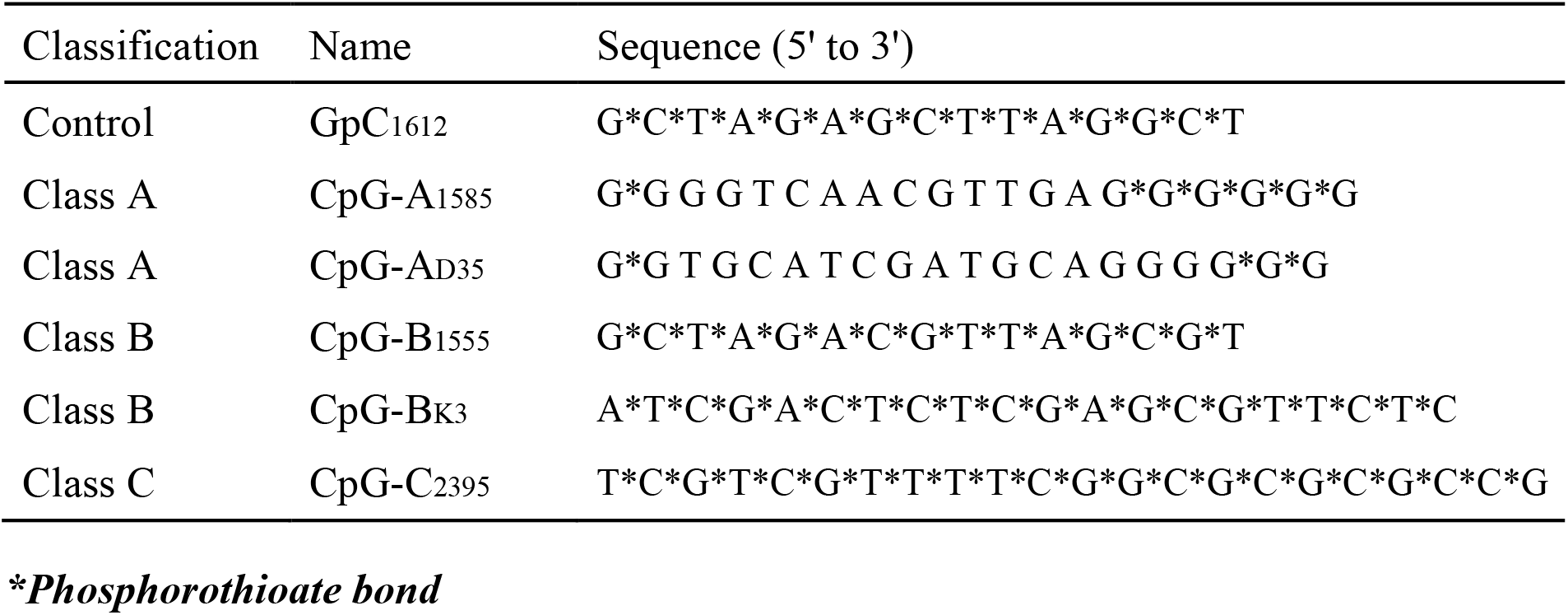
Nucleotide sequences of CpG-ODNs used in this study.

Seeded HD11 cells were incubated with 3 µM of each CpG-ODN for 24 h and subjected to cytotoxicity assays and quantitative reverse transcription-polymerase chain reaction (**qRT-PCR**) analysis. In addition, HD11 cells incubated with 3 µM of each ODN for 24 h, followed by stimulation with 500 ng/mL LPS for 2 h, were also assayed using the above methods.

### Cytotoxicity assay

The proportion of CpG-ODN and LPS-treated cells was observed using the inverted microscope IX71 (Olympus, Tokyo, Japan) and photographed using a DP70 camera (Olympus). Cell viability was assessed by the 3-(4,5-dimethylthiazol-2-yl)-2,5-diphenyl tetrazolium bromide (**MTT**) assay with some modifications (Ito et al., 2013; Hattori et al., 2021). Briefly, 300 µL complete DMEM containing 0.5 mg/mL MTT was added to the washed cells, followed by incubation for 4 h at 37 °C. Non-internalized MTT was then washed away, and the cells were lysed with 300 µL of 40 mM hydrochloric acid-isopropanol solution to release the MTT internalized by the viable cells. Next, 100 µL MTT solution was transferred to a 96-well plate (Thermo Fisher Scientific), and the MTT concentration was measured colorimetrically. Further, the cytotoxicity was estimated as CpG-ODN or CpG-ODN plus LPS addition upon PBS control by measuring the optical density at 570 nm. The effect of actinomycin D, a cytotoxic reagent (FUJIFILM Wako Pure Chemical Corporation, Osaka, Japan), was also examined to confirm the proportion and viability of HD11 cells used in this study.

### qRT-PCR analysis

Total RNA from cells treated with CpG-ODN and LPS was purified using the FastGene™ RNA Premium Kit (NIPPON Genetics Co., Ltd, Tokyo, Japan). cDNA was prepared by the reverse transcription of 100 μg total RNA per sample using SuperScript IV reverse transcriptase (Thermo Fisher Scientific). Equal volumes of cDNA were used for quantifying various cytokine immune response-related factor cDNAs via qRT-PCR using the StepOne real-time PCR system (Applied Biosystems, Foster City, California). qRT-PCR analysis was performed using KOD SYBR qPCR Mix (Toyobo Co. Ltd., Osaka, Japan) with specific primers. The nucleotide sequences of primers used in this reaction are listed in Table 2. qRT-PCR reactions were performed in duplicate. The relative expression levels of each target were calculated using the ΔΔCt method (Livak and Schmittgen, 2001) and normalized by the scores of β-actin.

**Table 2.**
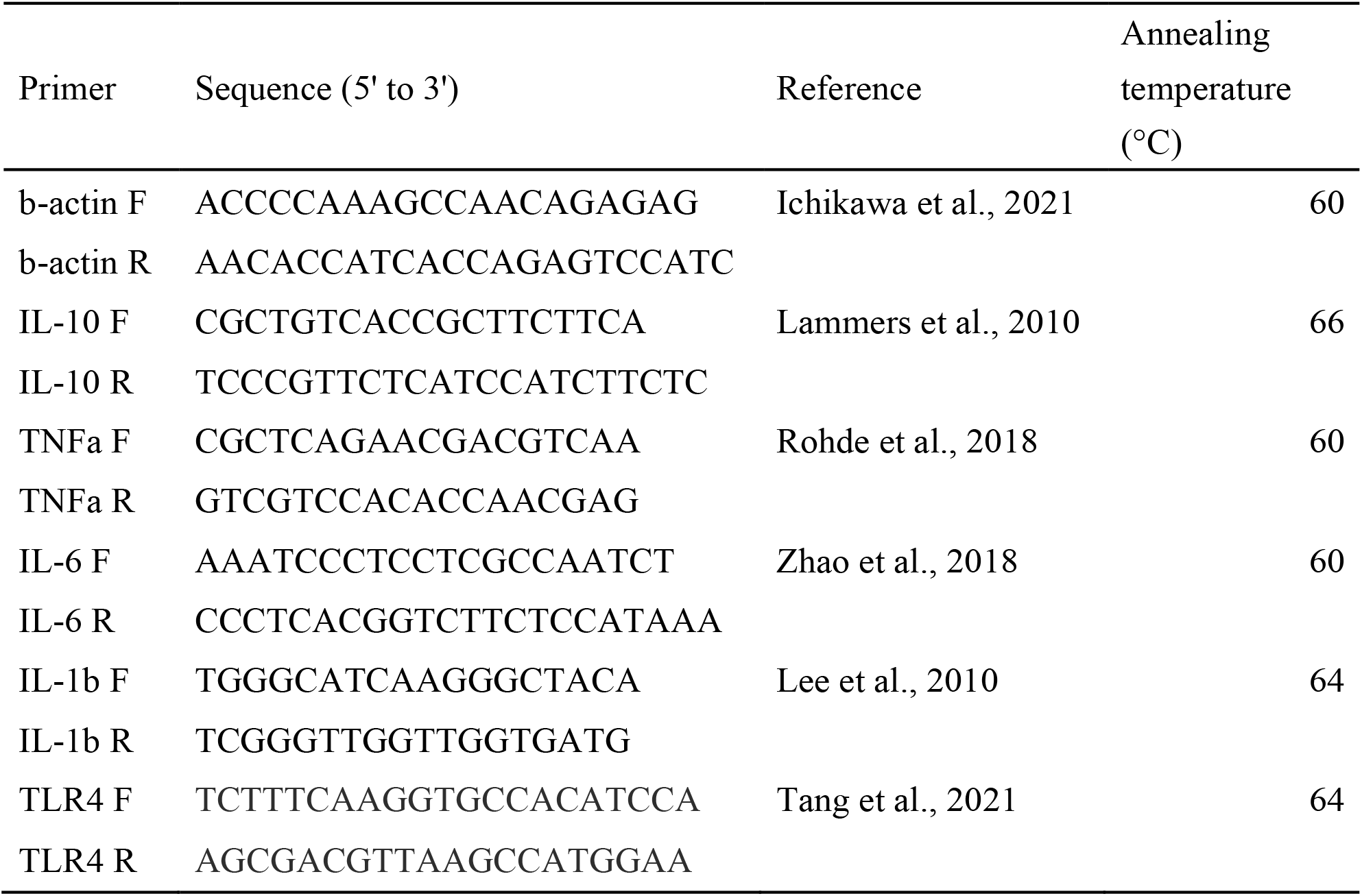
Nucleotide sequences of primers used in RT-qPCR.

### Statistical analysis

Statistical analyses were performed using GraphPad Prism8 (GraphPad Software Inc., LA Jolla, California). Data obtained from the cytotoxicity assay were analyzed using Dunnett’s test. Further, data obtained from qRT-PCR were analyzed using the one-way analysis of variance with the Tukey-Kramer post-hoc test.

## RESULTS

### Cytotoxicity of CpG-ODN in HD11 cells under Normal and Inflammatory conditions

We examined whether the viability of HD11 cells was affected by CpG-ODN using the IX71 inverted microscope and MTT assay. Microscopic observations showed that treatment with all CpG-ODN did not affect the proportion of HD11 cells under normal conditions. However, actinomycin D treatment reduced the proportion of cells under the same conditions (Figure 1A). The MTT assay showed that treatment with CpG-A_1585_, CpG-B_1555_, CpG-B_K3_, and CpG-C_2395_ significantly increased the cell proliferation index compared to only medium (Figure 1B), thereby exhibiting enhanced mitogenic activity. Thus, it is likely that CpG-ODN, except CpG-A_D35_, enhance mitogenic activity in HD11 cells under normal conditions.

**Figure 1.**
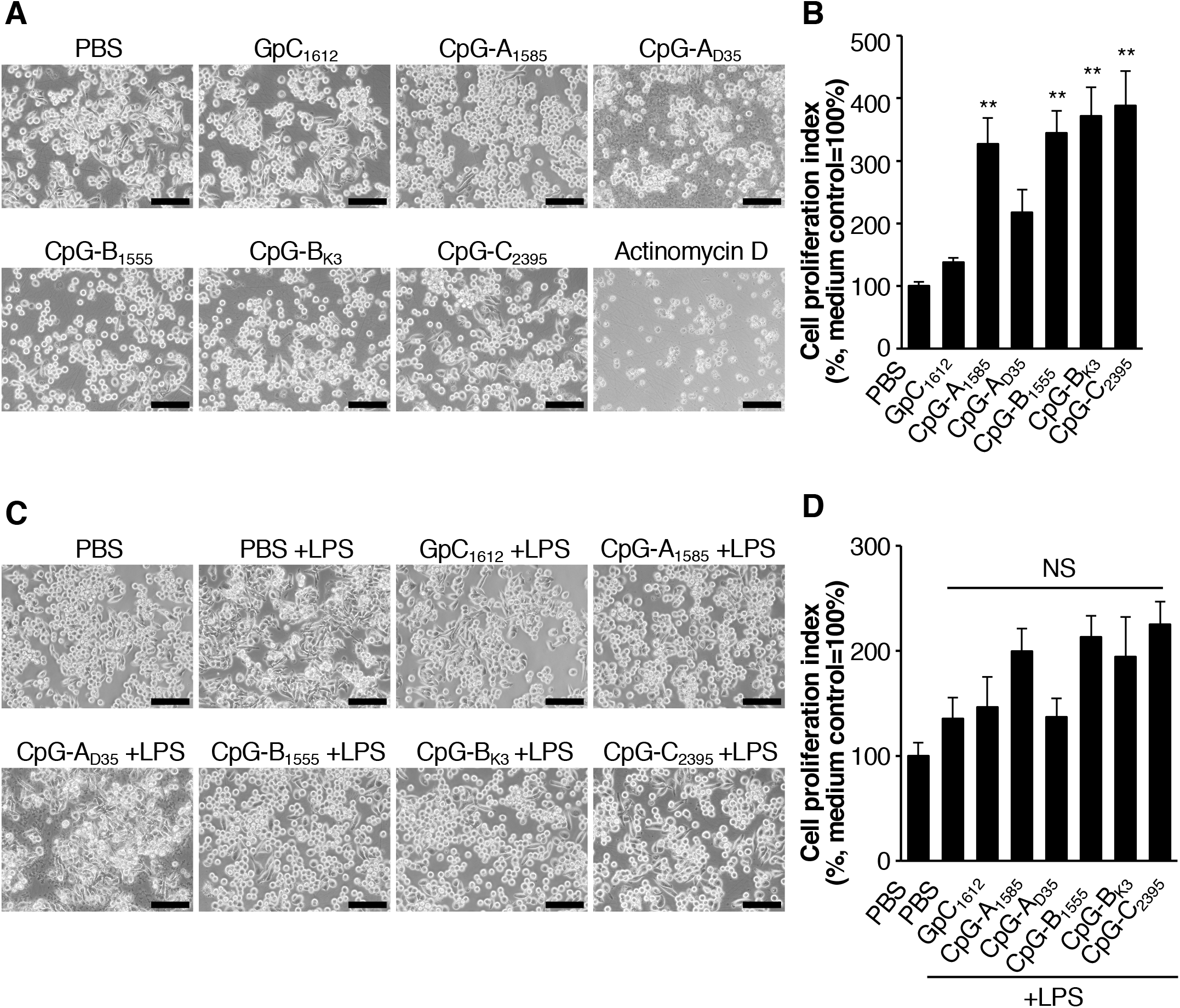
Evaluation of cellular injury and proliferation/viability. (A, D) Schematic diagrams of experimental design using oligodeoxynucleotides (**ODN**) without (A) or with (D) lipopolysaccharide (**LPS**). (B, E) Cellular morphology of HD11 cells treated with ODN (B) or stimulated by LPS after pretreatment with each ODN (E). Scale bars = 100 µm. (C, F) The results of 3-(4,5-dimethylthiazol-2-yl)-2,5-diphenyl tetrazolium bromide **(MTT)** assay using HD11 cells treated with ODN (C) or stimulated by LPS after pretreatment with each ODN (F). Error bars indicate standard error of the mean (**SEM**) of the cell proliferation index calculated from three independent experiments (n = 3). The statistical significance was evaluated by Dunnett’s test compared with the HD11 treated with PBS (C) or stimulated by LPS after pretreatment with PBS (F) (\*\**P* < 0.01). NS means not significant.

Next, the same analyses were conducted using HD11 cells under inflammatory conditions. The cell proportion did not change under these conditions (Figure 1C). However, contrary to the results of the MTT assay under normal conditions, pretreatment with all CpG-ODN did not change the cell proliferation index compared to LPS alone (Figure 1D). Therefore, all CpG-ODN used in this study were non-toxic to HD11 cells under both normal and inflammatory conditions.

### Comparison of CpG-ODN on induction of IL-10 mRNA expression in HD11 cells

To screen for CpG-ODN that strongly induce anti-inflammatory cytokines in HD11 cells, we examined the effects of CpG-A_1585_, CpG-A_D35_, CpG-B_1555_, CpG-B_K3_, and CpG-C_2395_ on *IL-10* mRNA expression in HD11 cells under normal and inflammatory conditions. Treatment with CpG-B_1555_, CpG-B_K3_, and CpG-C_2395_ significantly induced *IL-10* mRNA expression compared to other ODN and the PBS control under normal conditions, with the highest induction level in CpG-C_2395_ (Figure 2A). In addition, pretreatment with CpG-C_2395_ resulted in the highest induction of *IL-10* mRNA expression under LPS-induced inflammatory conditions. However, similar results were not observed in HD11 cells pretreated with CpG-B_1555_ and CpG-B_K3_ plus LPS (Figure 2B). Thus, CpG-C_2395_ strongly induces IL-10 in HD11 cells under both conditions. Therefore, we selected CpG-C_2395_ as an effective inducer of IL-10 and used it for subsequent functional analyses, focusing on the regulation of inflammatory responses.

**Figure 2.**
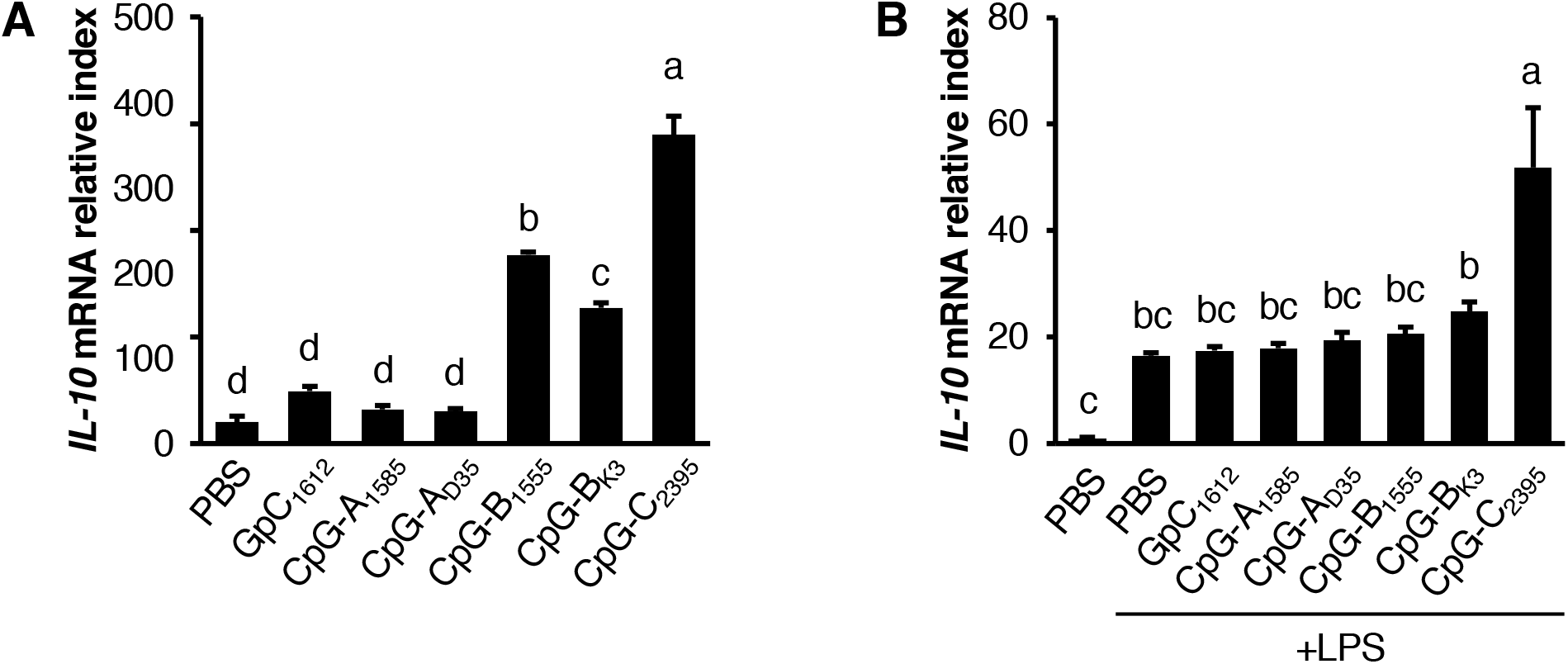
Expression analysis of IL-10 using quantitative reverse transcription-PCR (**qRT-PCR**). (A) Expression analysis of HD11 under the single-stimulus condition. (B) Expression analysis of HD11 stimulated by LPS after pretreatment with each ODN. The ΔΔCt method was used for the calculation of relative expression levels. The levels of *IL-10* were normalized to β-actin mRNA expression. Error bars indicate SEM of the relative expression levels during three independent experiments (n = 3). Statistical significance was evaluated by the Tukey’s test. The different alphabets represent significant differences (*P* < 0.05).

### Effects of CpG-C_2395_ treatment on LPS-induced Inflammatory responses in HD11 cells

Finally, we evaluated the effects of CpG-C_2395_ pretreatment on LPS-induced inflammatory responses. TNF-α, IL-1β, and IL-6 were selected as candidates for pro-inflammatory cytokines. The mRNA expression levels of *TNF-α, IL-1β*, and *IL-6* remarkably increased in cells treated with LPS, but pretreatment with CpG-C_2395_ significantly inhibited these cytokines. Additionally, the mRNA expression level of the LPS-induced toll-like receptor (***TLR***) *4*, a pattern recognition receptor that specifically recognizes LPS, was also significantly inhibited by CpG-C_2395_. These results suggest that treatment with CpG-C_2395_, before inflammatory responses are induced by LPS, contributes to the inhibition of inflammation in HD11 cells.

## DISCUSSION

In the current study, we aimed to characterize the anti-inflammatory responses of a chick macrophage cell line, HD11, by comparative analysis using several CpG-ODN. A class C CpG-ODN, CpG-C_2395_, directly upregulated *IL-10* expression and reduced LPS-induced inflammation. To the best of our knowledge, this is the first report that sheds light on the anti-inflammatory responses in chick macrophages using a comparative analysis of three major classes of CpG-ODN.

Single-stimulus of each ODN in HD11 cells showed that CpG-Bs and CpG-C_2395_ could induce *IL-10* expression (Figure 2A) without cell injury (Figures 1B and 1C). Generally, CpG-As stimulate plasmacytoid dendritic cells (**pDCs**) and induce the expression of interferon-α. CpG-Bs mainly target B cells and induce immunostimulatory responses. CpG-Cs possess intermediate structural features between CpG-As and CpG-Bs and can stimulate both pDCs and B cells (Hanagata, 2012). CpG-Bs can directly stimulate mammalian macrophages and induce IL-10 expression (Yi et al., 2002; Boonstra et al., 2006). Moreover, CpG-A_1585_ can also induce anti-inflammatory responses against septic shock in mice (Yamamoto et al., 2017). While a single-stimulus CpG-Bs or CpG-Cs can induce *IL-10* expression, CpG-A_1585_ can stimulate cellular proliferation/viability without upregulating *IL-10* expression. As the functions of CpG-As in chickens are poorly understood, further studies are needed.

Pretreatment with each ODN before LPS stimulation showed that only CpG-C_2395_ promoted the expression of *IL-10* under inflammatory conditions (Figure 2B). Additionally, pretreatment with CpG-C_2395_ downregulated several inflammation-related genes without cell injury (Figures 1E, 1F, and 3). Thus, CpG-C_2395_ could induce anti-inflammatory responses following LPS stimulation. Previous comparative studies using some classes of CpG-ODN have demonstrated the efficiency of CpG-Bs against antiviral or bacterial responses in chickens (Dar et al., 2009; Barjesteh et al., 2014). Additionally, CpG-Bs were also identified upon administering CpG-ODN as a vaccine adjuvant into chickens (Wang et al., 2009). Our findings suggested that anti-inflammatory responses were strongly induced by treatment with CpG-Cs but not with CpG-Bs.

**Figure 3.**
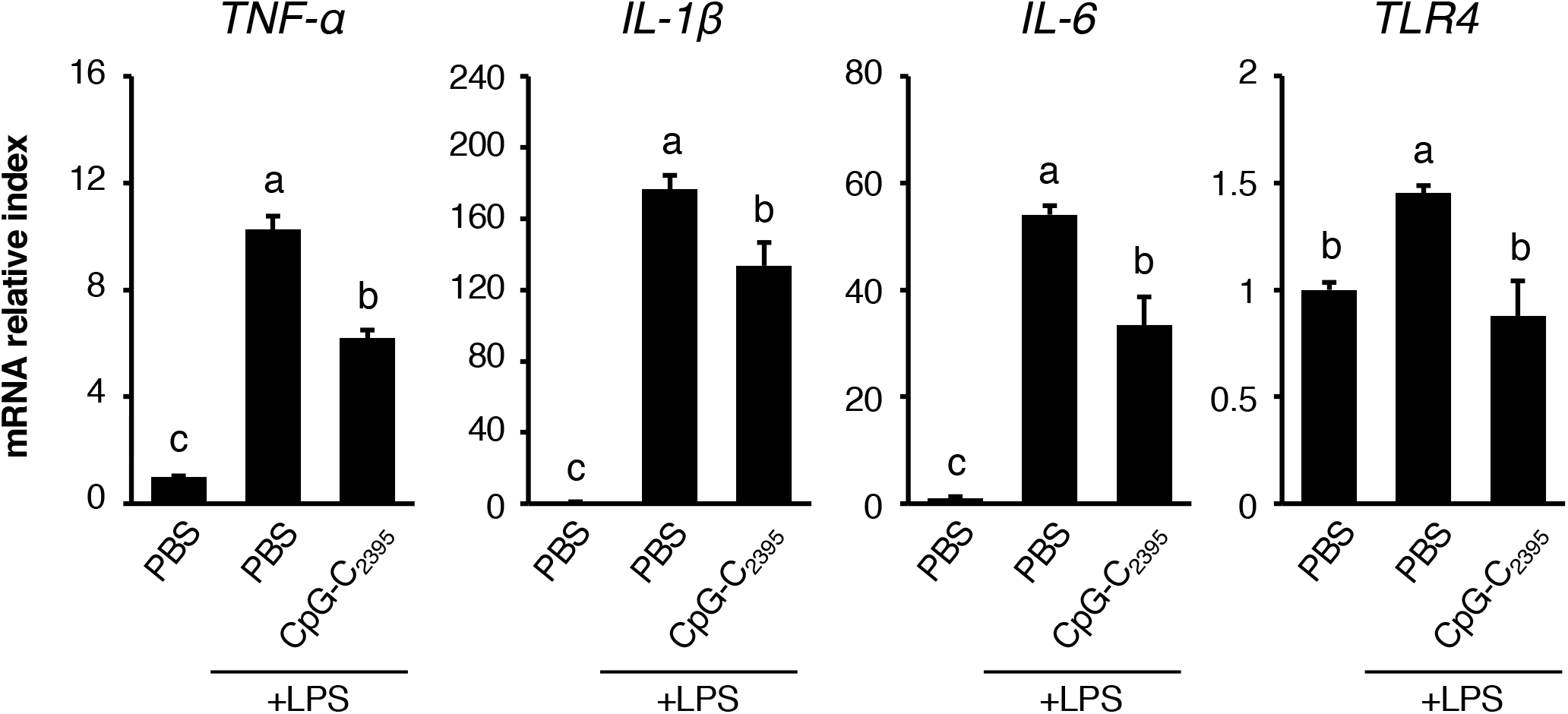
Expression analysis of the inflammation related genes using qRT-PCR. The ΔΔCt method was used for the calculation of relative expression levels. The levels of each gene were normalized to the expression of β-actin mRNA. Error bars indicate SEM of the relative expression levels during three independent experiments (n = 3). Statistical significance was evaluated by Tukey’s test. The different alphabets represent significant differences (*P* < 0.05).

Avian cells possess characteristic mechanisms for recognizing CpG-ODN. In mammals, CpG-ODN is recognized by TLR9 (Hemmi et al., 2000). However, birds have evolutionally lost TLR9, and TLR21 acts as a functional homolog (Brownlie et al., 2009; Keestra et al., 2010). Interestingly, although mammalian TLR9 and avian TLR21 are functionally identical in recognizing CpG-ODN, they have minimal sequence similarity and different CpG-ODN sequence recognition profiles (Xie et al., 2003; Keestra et al., 2010; Chuang et al., 2020). Additionally, TLR15, a Toll-like receptor unique to birds, also contributes to CpG-ODN recognition (Ciraci and Lamont, 2011). Therefore, to use CpG-ODN as an anti-infective agent in the poultry industry, basic knowledge of the immunostimulatory features of CpG-ODN against avian cells, not mammalian cells, is required. Our findings provide a new perspective on the superiority of CpG-Cs for suppressing diseases caused by the excessive expression of pro-inflammatory cytokines, such as sepsis and avian influenza, in chickens.

In summary, we examined the effect of pretreatment with each CpG-ODN against anti-inflammatory responses triggered by LPS in chick macrophages. However, we have not yet evaluated cell-cell interactions and functions against infection following pretreatment with CpG-ODN. Hence, further studies are needed to evaluate the effects of CpG-ODN in vivo and to apply the findings of this study to the poultry industry. Recently, an oral delivery system of CpG-ODN using nanocapsules has been established in mice (Wang et al., 2015). For applying our findings to the poultry industry, a simple method of administering CpG-ODN, such as nanocapsules, would be helpful.

## ACKNOWLEDGMENTS

This research was supported in part by Japan Society for the Promotion of Science KAKENHI Grant Number 21K14997. We would also like to thank Editage (http://www.editage.jp) for their help in English language editing.

## DECLARATIONS OF INTEREST

None.

## Notes

### Competing Interest Statement

The authors have declared no competing interest.

